# Non-invasive canine electroencephalography (EEG): a systematic review

**DOI:** 10.1101/2023.08.14.552730

**Authors:** Akash Kulgod, Dirk van der Linden, Lucas G S França, Melody Jackson, Anna Zamansky

## Abstract

The emerging field of canine cognitive neuroscience uses neuroimaging tools such as electroencephalography (EEG) and functional magnetic resonance imaging (fMRI) to map the cognitive processes of dogs to neural substrates in their brain. Within the past decade, the non-invasive use of EEG has provided real-time, accessible, and portable neuroimaging insight into canine cognitive processes. To promote systematization and create an overview of framings, methods and findings for future work, we provide a systematic review of non-invasive canine EEG studies (N=22), dissecting their study makeup, technical setup, and analysis frameworks and highlighting emerging trends. We further propose new directions of development, such as the standardization of data structures and integrating predictive modeling with descriptive statistical approaches. Our review ends by underscoring the advances and advantages of EEG-based canine cognitive neuroscience and the potential for accessible canine neuroimaging to inform both fundamental sciences as well as practical applications for cognitive neuroscience, working dogs, and human-canine interactions.

## Introduction

The multidisciplinary field of cognitive neuroscience is a synthesis of cognitive psychology and neuroscience, aiming at mapping “elementary cognitive functions onto specific neuronal systems” [3, pg. 613]. Through the deployment of techniques such as the electroencephalogram (EEG) [98] and functional magnetic resonance imaging (fMRI) [29,105], neuroscientists are able to perform empirical and quantitative analyses of cognitive processes. For a comprehensive review of the history, methods, and current frameworks of cognitive neuroscience, readers are directed towards Gazzaniga, Ivry and Mungun’s 2019 treatment of the field [30]. Another multidisciplinary field is canine science, combining disciplines including evolution, genetics, cognition, ethology, physiology, comparative medicine, and ecology [14, 15, 69, 74]. Investigating the recent surge of interest in the scientific study of the domestic dog, Aria and colleagues [7] found a sixfold increase in the number of studies in canine cognition and behaviour between the years 2006 and 2018, as compared to the preceding period of 1993 to 2005. This interest extends beyond purely veterinary, pharmaceutical, and basic neuroscience paradigms. Recent studies in canine science span such varied topics as genetics [32], evolutionary neuroscience [36, 37] and intelligence [6], as well as investigations into models of epilepsy [68], aging [51, 73] and dementia [95]. Recent developments, such as the rise of open-science multi-team initiatives such as the ManyDogs Project, which aims to investigate behavioral traits across multiple centers and populations of dogs [71], and the Working Dog Project, which focuses on improving genetic selection strategies in dog breeding [20], herald the emerging trend of collaborative consortiums to tackle fundamental questions in the field.

Berns et al. pioneered the use of fMRI in awake, non-restrained dogs in 2012 [8], and in the following year, Kujala and colleagues were the first to successfully deploy non-invasive EEG with non-sedated dogs [61]. Further developments included the investigation of a range of cognitive processes and their neural underpinnings such as executive functioning [24], visual [25, 61], auditory [5, 17], and olfactory [50] processing, social cognition [23], learning [88] and sleep [19, 56]. While the field of canine fMRI has received increasing scientific attention, non-invasive canine EEG has eluded similar treatment. This is despite its noticeable strengths, including high temporal resolution, accessibility, and real-world applications. For these reasons, we offer a consolidation of the state of EEG-based canine cognitive neuroscience, providing an overview of the key conceptual framings, methodological approaches, and findings.

To that end, this systematic review contributes the following:

**An identification and mapping of 22 non-invasive canine EEG studies** based on a thorough literature review and an tailored database query paired with appropriate exclusion/inclusion criteria.

**A systematic analysis of these studies** dissecting their research question and participant make-up, technical setups deployed, dataset properties, and analytical frameworks and findings.

**A critical discussion on future avenues for non-invasive canine EEG** identifying promising questions that can be pursued, ideal data practices, integration with other research sub-fields and beneficial methodological and analytical refinements.

## Methods

### Data sources and search query construction

No literature reviews, surveys or meta-analysis of EEG in dogs were available as a starting point for this review. To begin, we conducted an informal search using Google Scholar in February 2023, searching for ‘EEG in dogs’ and read through a number of obvious candidates ([10, 60, 61, 66, 70, 99] [17, 54–56]) to identify commonalities. Of these papers, we noted that publishers included PLOS One, Springer, Elsevier, and the Royal Society, while we would also expect papers published in IEEE and ACM venues to be relevant. This allowed us to minimize necessary redundancy in data sources by employing each publisher’s own search engine, as *Scopus* indexed all early identified papers, and papers by expected publishers.

After reading the initial set of studies, we made the following assumptions to aid in the construction of our search query, to ensure as broad a coverage as possible:

- Dogs are referred to interchangeably as ‘dog*’ or ‘canine*’, even if articles do not necessarily include the full scope of the canis family; this required an additional exclusion criterion if papers include e.g., *Canis lupus* rather than just *Canis lupus familiaris*.
- EEG is referred to both as ‘EEG’ and ‘Electroencephalography’ or more vaguely hinted at with terms such as ‘brain signals’ or ‘neural processes’ in paper titles, meaning we needed to search through abstracts as well.
- There is no consistent (from title) indication of whether the used EEG technique was invasive or non-invasive, this required liberal inclusion criteria to include any relevant study and stringent exclusion criteria to manually filter any invasive EEG study.
- The first non-invasive, non-sedated canine EEG study could be clearly identified as taking place in 2013 [61] so the year 2010 allowed for all relevant studies to be included.

Based on the above assumptions and reading of the initial papers, we constructed this search query optimized for the SCOPUS database in March 2023:

(TITLE ((canine* OR dog*)) AND TITLE-ABS-KEY ((eeg OR erp OR electroencephalography))) AND PUBYEAR > 2010 AND PUBYEAR < 2024 AND (LIMIT-TO (LANGUAGE, “English”))

The search query resulted in 205 articles in the SCOPUS database.

### Study selection

We first *liberally* applied the inclusion criterion, followed by a set of three exclusion criteria – see Fig. 1 for an overview of the entire process and interim study numbers. The inclusion criterion was:

**Figure 1:**
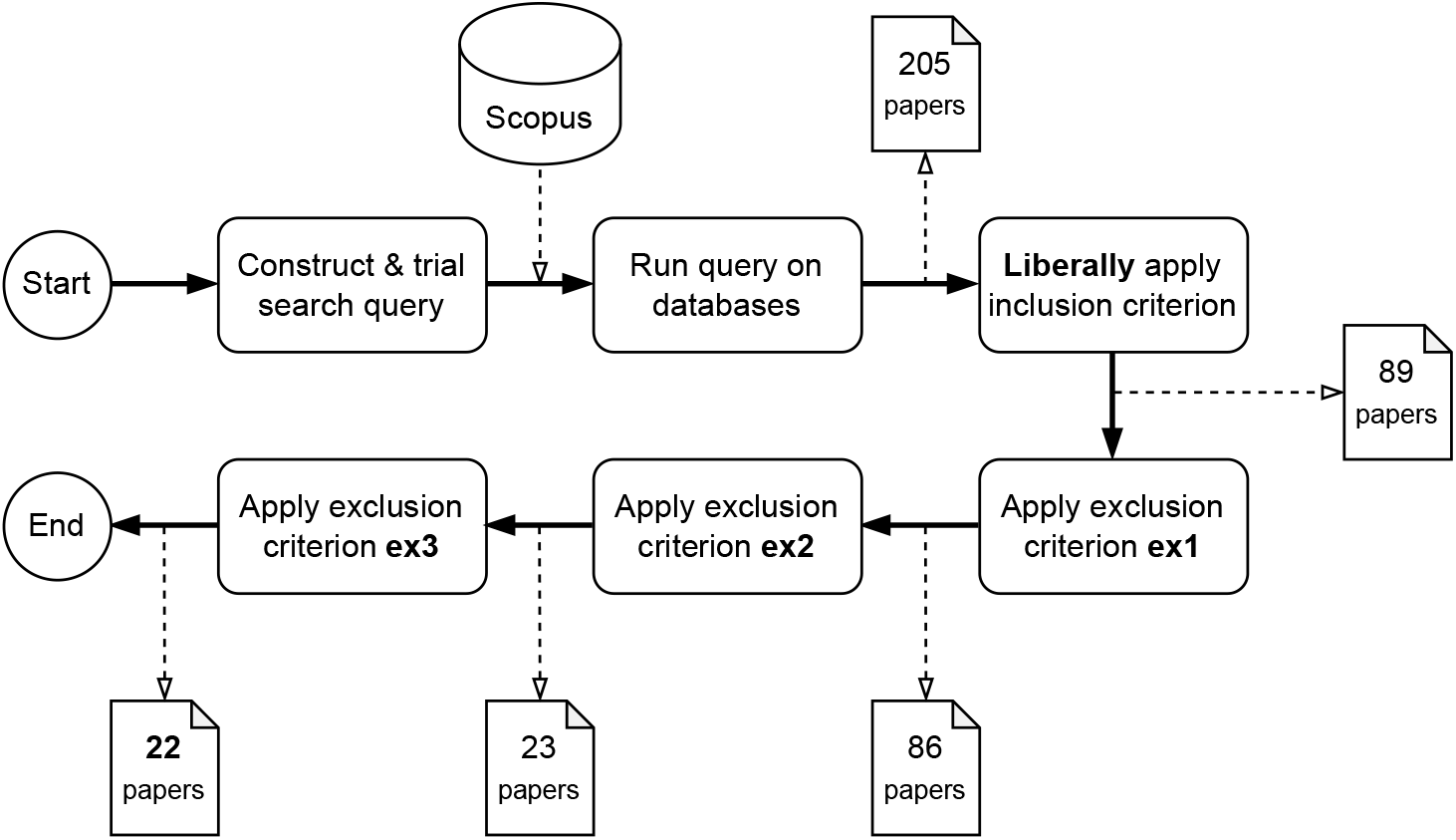
Flowchart of the study selection process and included papers at each step of the process.

**inc1** Studies applying electroencephalography to dogs (*canis lupus familiaris*) The following exclusion criteria were used to refine the selection:

**ex1** Studies of other canine species (e.g., wolves, jackals, coyotes) – we built in this potentially redundant exclusion criteria due to the polysemous use of ‘canine’ in some literature, sometimes referring to its actual meaning of the *Caninae* subfamily, but at other times used as a synonym for dogs only.

**ex2** Studies employing invasive applications of EEG – we define this as the physical ontology of the subject remaining unviolated, i.e. the epidermis of the subject dog is not pierced or excised. This excluded any study using intracranial EEG (iEEG) [83] as well as sub-dermal EEG that uses needle electrodes, a technique first used by Pellegrino and Sica (2004) [84] in a veterinary context as well as cognition studies such as by Howell and colleagues. [42]

**ex3** Studies employing anaesthesia or other form of sedation – we do include studies of naturally sleeping dogs, which, indeed, form the majority of the studies conducted in this category

One author applied the inclusion and exclusion criteria over the total set of 205 papers leading to a final selection of 22 papers. To ensure consistent application of the criteria, another author independently coding a randomly selected 10% subset of the papers. Inter-rater reliability analysis indicated substantial agreement between authors on application of the inclusion criterion (Cohen’s *κ*=0.69) criteria, as well as the exclusion criteria (resp. *κ*=0.69, 1.00, 1.00 for ex1, ex2, and ex3), leading to the same set of selected papers. We found that the validating author labeled more critically, which led to some discussion as to whether retrospective studies were to be included; but effectively these had no impact on the actual selected publications as most were ruled out by both authors on grounds of the exclusion criteria.

### Data extraction and analysis

We dissect the obtained studies according to the different workflow stages of a scientific study: research question and participants, data acquisition, and analysis/findings, respectively. This process was done in entirety manually by one author. After providing an overview of the 22 studies, we break them down along the following three dimensions -

- Setups deployed to collect data: electrode types, montages, and amplifiers.
- Datasets collected in each of the papers: e.g., availability and quantity of data.
- Analysis frameworks and findings: e.g., pre-processing pipelines (where relevant), and findings.

For each of these steps, we highlight and synthesize common themes and approaches and use this to provide a consolidated outlook on the field and suggestions for future work.

## Results

### Overview

The selected works are presented in Table 1. The dimensions we use and the overall trends of the studies were as follows :

**Table 1:**
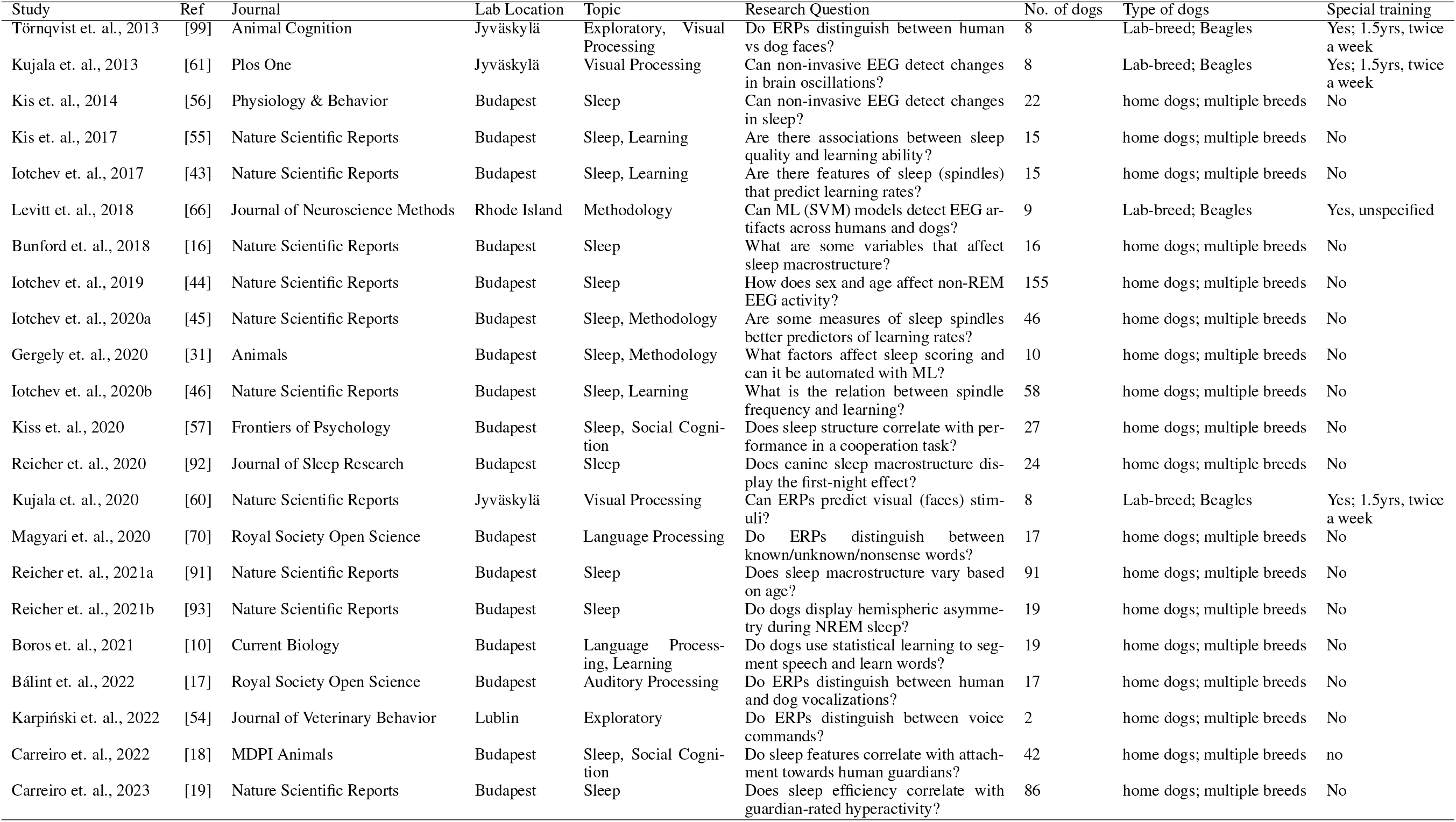
Overview of the 22 reviewed studies.

- *Topic investigated*. We label each study with one or more of the following categories -
  - visual processing : perception, discrimination or interpretation of images (n=3)
  - auditory processing : perception, discrimination or interpretation of sounds (n=1)
  - language processing : perception and comprehension of speech (n=3)
  - learning : associative learning, memory, and problem-solving (n=4)
  - emotion : interpretation of emotionally coded stimuli (n=2)
  - social cognition : behavior with con-specifics or humans (n=2)
  - sleep : stages and occurrence of patterns during sleep (n=14)
  - methodology : approaches to data collection and analysis such as automation or application of machine learning (n=3)
- *Specific research question*. We briefly state the research question/hypothesis investigated in each study.
- *Number of dogs*. The number of participants in each study ranged from 2 [54] - 155 [44].
- *Type of dogs*. Here we refer to home/laboratory dogs, and their breed (specific/multiple).
- *Dog Training*. In this category, we describe the nature of training provided to the dogs for data collection in the study, distinguishing between studies that employed special training and those that did not. We use special training to refer to a dedicated training process consisting of multiple preparatory sessions, such as habituation to the equipment, conducted prior to actual data collection.

Overall, four different research centers in North America and Europe have conducted non-invasive canine EEG studies, with the majority of them coming from Eötvös Loránd University in Budapest, Hungary. The majority of studies (n=14) recorded EEG from sleeping dogs, taking advantage of the ease of recording higher quality data for longer periods and the presence of well-defined cross-species EEG structures associated with sleep stages and patterns.

The majority of dogs were companion (home) dogs whose guardians were recruited using surveys, and consisted of a diverse group of pure and mixed breeds across ages in both male and female animals. Four studies [60, 61, 66, 99] record from purpose-breed laboratory Beagles and only these dogs underwent extensive training for EEG recording purposes, whereas companion dogs were habituated to equipment on the same day as recorded sessions.

### EEG Setups

To facilitate a systematic and rigorous comparison of studies, we compiled Table 2 outlining the six different canine EEG setups used by reviewed studies. Some important dimensions of the setups are a) electrodes type and attachment, which includes the type of electrodes, method of attachment, whether fur was shaved, and the impedance of signals (a measure of conductivity at the electrode-skin interface); b) the number and montage of electrodes, which refers to the number of recording channels and the specific arrangement of electrodes; and c) the amplifier model and sampling rate.

**Table 2:**
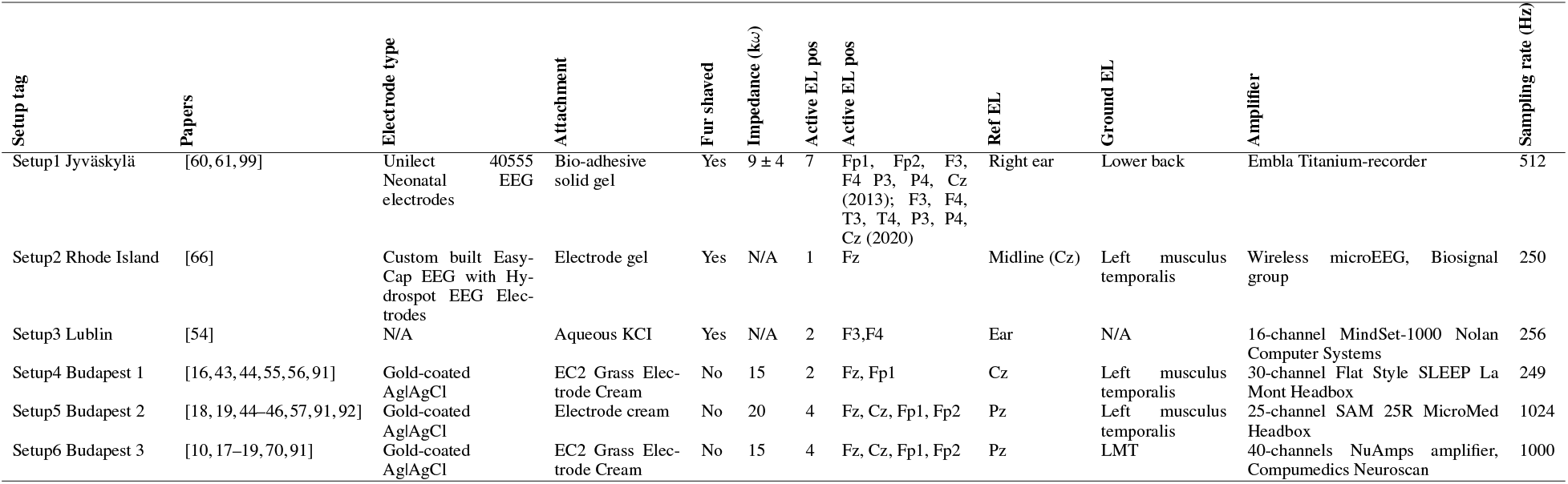
Overview of the six EEG setups used by the reviewed studies.

### Electrodes type and attachment

Electrodes are made of different conductive materials and are generally categorized along two binaries -gel vs dry, and active vs passive. The benefit of different electrode types are still debated [39, 106], however, with regards to our review, all studies used passive gel electrodes secured with surgical tape or electrode cream. Four studies shaved the fur of participant dogs [60, 61, 66, 99], with the goal of ensuring higher quality data by decreasing impedance, however studies that did not shave fur have achieved comparable levels of impedance. Higher impedance values result in a lower signal-to-noise ratio (SNR), although optimal electrode impedances vary relative to an amplifier’s input impedance [85] (but see also Kappenman and Luck [52] for a review). All studies reported keeping impedance values below 20 kΩ.

### Electrodes number and position

Canines have smaller brains than humans and subsequent less space available for electrodes. The reviewed studies used between one to seven electrodes, along with a ground and reference. In comparison, human EEG studies typically vary between using 4-256 active channels [65]. However, it is worth noting that dogs have far fewer cortical neurons than humans, about 500 million [48] compared to 16 billion [38] in humans -a ratio of 1:32. Arguably, this means that a 4 channel recording in dogs is equivalent to a 128 ‘high density’ recording in humans.

Electrode montages in EEG research refers to the “logical, orderly arrangements of electroencephalographic derivations or channels that are created to display activity over the entire head and to provide lateralizing and localizing information.” [1]. Human EEG research generally uses the 10-20 montage system, formalized in 1957-58 by Herbert Jasper [49], to standardize electrode montages and ensure replicability across studies. Canine EEG has traditionally borrowed from the human 10-20 system and the reviewed studies use similar derivations, although setups differed from each other in the montages used and authors differed in the labels used for specific montage positions. For an outlook of the montages used in the reviewed studies, see Figure 2A notable challenge with standardizing electrode montages in dogs is the remarkable variance in head shape and size across canine breeds. One way to solve this challenge is to record only from dogs of the same breed and avoid it all altogether, which was the case with the studies recording only from lab-bred Beagles. [60, 61, 66, 99]. Another approach is to use the relative distance between breed-invariant anatomical markers followed by the Budapest setups [56, 70].

**Figure 2:**
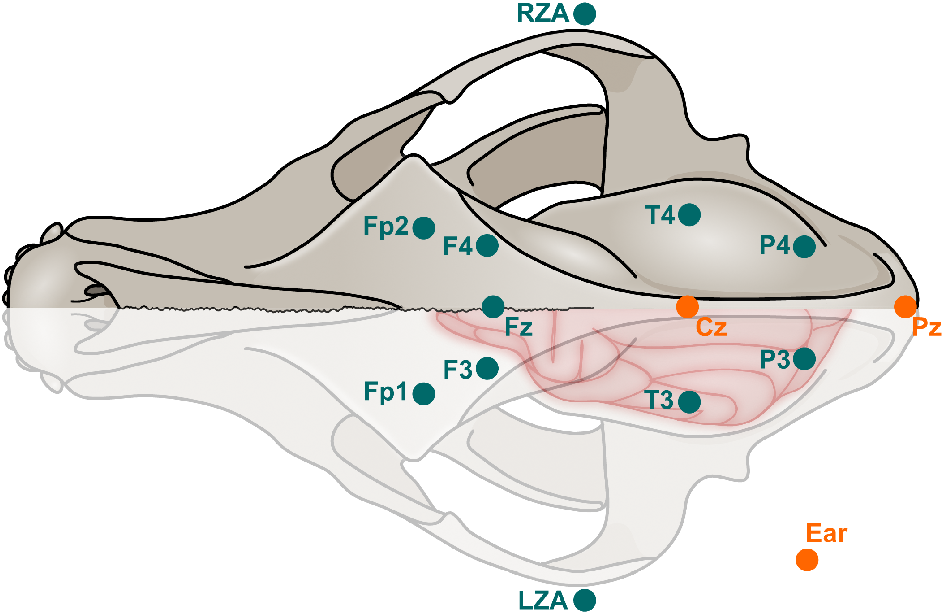
Different electrode montages used by the six technical setups. Electrode labels follow from the human 10-20 system, where letters indicate the lobe — Fp=pre-frontal, F=frontal, P=parietal, T=temporal, and C=center. The left and right zygomatic arch are also depicted as LZA and RZA. Odd numbers refer to electrode placement on the left side and even numbers indicate placement on the right side of the brain. Reference electrode positions are highlighted in orange. It is worth pointing out that large variance amongst canine breeds means the figure is not representative of all canine individuals.

Traditional electrode montages on humans when translated to dogs might suffer from a higher rate of muscular artifacts due to the presence of a muscular scalp in dogs, which motivated the authors of the Budapest setups to place electrodes on the anteroposterior midline, or sagittal crest, of the canine skull (Fz, Cz, and Pz respectively), as it is a bony ridge that minimizes muscular artifacts. These setups also used either 1 or 2 electrodes close to the eyes (Fp1 and Fp2) to measure electrooculography (EOG) signals. EOG signals are useful for understanding eye activity, such as blinks, which can have an effect on recordings from other electrodes. Jyväskylä setups differed in their approach by positioning electrodes laterally - on the frontal cortex F3 and F4, on the posterior cortex P3 and P4, and the temporal cortex T3 and T4 (the latter only in the 2020 study [60]).

Finally, the position of the reference electrode plays a key role in recordings as EEG data is derivational and signals from any electrode are meaningful only in respect to a reference electrode (or electrodes). In the Budapest setups, either the center of the sagittal crest (Cz) or the bony ridge of the occipital protuberance, or occiput, (Pz) was used as a reference, given its relative distance from both neural and muscular activity. The Rhode Island setup also used a Cz reference. Meanwhile, Jyväskylä setups used an electrode placed on the ear as the reference, as did the sole Lublin study. Figure 2 gives an overview of different potential electrode montages used across the selected studies.

### Amplifier model and sampling rate

Various factors determine amplifier performance including sampling rate, input range, amplifier impedance, band-width, and the common mode removal ratio (CMRR). A detailed discussion of these factors is outside the scope of this review (readers are directed towards the 2023 review of EEG systems by Niso and colleagues [79] for a discussion). One feature worth highlighting is the sampling rate which refers to the number of samples that are digitally acquired per second (measured in Hz). Higher sampling rates are useful primarily to measure higher frequency brain activity, given by the Nyquist-Shannon theorem [103] that states that for any periodic signal of a given frequency, a sampling rate higher than two times the frequency of the signal is needed to accurately detect its presence. Sampling rates of amplifiers used ranged from 250-1024 Hz. As all studies focused analysis on bandwidths between 0-50Hz, all amplifiers had a sufficient sampling rate.

### Datasets

The six different technical setups described in the prior section were used to acquire 18 datasets, summarized in Table 3. The discrepancy between number of papers and datasets is because some papers developed novel methodological approaches [43] or refinements [31, 45] using older datasets, or deploying a different analytical framework on the same dataset [61]. Many studies use a combination/amalgamation of different data sources so the datasets presented here should not be assumed to be well-demarcated and unique. Moreover, the overall number of dogs is difficult to establish, as we observed a considerable overlap of participants between studies and datasets, with the same dogs being recorded multiple times and the same datasets being reused in different papers. Datasets were categorized into either epoch or continuous paradigms, based on which, the number of events/minutes of data per dog and per dataset was calculated. All but one [66] of the awake dogs datasets were epoched and all sleep datasets were continuous. Finally, we noted the availability of the datasets. No datasets were freely available in their entirety and six datasets were partially available, either requiring a) additional author authorization on a data repository; b) an email request; c) missing important metadata; d) missing raw data, i.e. containing only processed data.

**Table 3:** Overview of the 18 datasets created by non-invasive canine EEG studies.

| Name | Papers | No. of dogs | Unique dogs? | Type | Conditions | Events OR sessions per dog | Duration of recording per dog (min) | Total events OR duration of data | Open source? |
| --- | --- | --- | --- | --- | --- | --- | --- | --- | --- |
| Faces1 | [61, 99] | 8 | Yes | Epoch, Continuous | 2 | 240 | 80 | 976 | No |
| Sleep_Exploratory | [56] | 22 | Yes | Continuous | 1 | 1 | 180 | 3960 | No |
| Sleep_Exploratory2 | [56] | 7 | No | Continuous | 2 | 2 | 360 | 2520 | No |
| Sleep_Learning1 | [55] | 15 | Unsure | Continuous | 2 | 2 | 360 | 5400 | No |
| Auto_Artefacts | [66] | 9 | Yes | Continuous | 1 | 90 | N/A | N/A | No |
| Sleep_Activity | [16] | 16 | Unsure | Continuous | 2 | 2 | 540 | 8640 | No |
| Sleep_Variation | [44] | 155 | Unsure | Continuous | 1 | 1 | 180 | 27900 | No |
| Sleep_Aging | [45] | 58 | Unsure | Continuous | 2 | 2 | N/A | N/A | No |
| Sleep_Individual | [57] | 27 | Unsure | Continuous | 1 | 1 | 180 | 4860 | No |
| Sleep_Adapt | [92] | 24 | Unsure | Continuous | 3 | 2 | 180 | 4320 | No |
| Faces2 | [60] | 8 | Unsure | Epoch | 8 | 1200 | N/A | 8000 | Partially A |
| Phoentics | [70] | 17 | Unsure | Epoch | 3 | 240 | N/A | 4080 | Partially B |
| Sleep_Development | [91] | 91 | Partial | Continuous | 1 | 3 | 540 | 49140 | No |
| Statistical_Learning | [10] | 19 | Unsure | Epoch | 4 | 320 | 12 | 6080 | Partially B |
| Vocalizations | [17] | 17 | Unsure | Epoch | 4 | 192-384 | N/A | N/A | Partially C |
| Pilot_Exploratory | [54] | 2 | Yes | Epoch | 2 | 2 | N/A | N/A | No |
| Sleep_Attachment | [18] | 43 | Unsure | Continuous | 1 | 1 | 1-3h | N/A | Partially D |
| Sleep_Hyperactivity | [19] | 86 | Unsure | Continuous | 1 | 1 | 1.5-3h | N/A | Partially D |

### Analysis and Findings

We review and consolidate awake and asleep dog studies separately, given their different frameworks, workflows, and results.

### Preprocessing

Preprocessing pipelines in EEG data analysis involve several steps to clean and prepare the data. One of these steps is artifact removal that aims to identify and remove segments of the signal containing unwanted noise or corruption. These artifacts can originate from various non-neural sources, such as eye blinks, muscle activity, or external interference [76]. Artifact removal can be performed using automated methods that apply predefined thresholds, such as amplitude criteria, to detect and exclude segments with excessive noise. Alternatively, manual inspection techniques like independent component analysis (ICA) or video monitoring can be employed. The choice between manual and automated detection methods is a topic of ongoing debate in the field, as each approach has its strengths and limitations. [47]

Reviewed studies differed in their application of pre-processing techniques, which have potential important ramifications for the analysis and validity of their respective findings. For example, Magyari et al. (2020) [70] tested two different artifact removal procedures - a multi-level method combining quantitative and qualitative steps, as well as a single-step approach using filtering and amplitude-based artifact removal. The multi-level approach consisted of automated amplitude-based rejection, manual video coding of movement and manual inspection of EEG data. The single-step consisted only of automated artifact removal and filtering. The results showed similar condition differences between the two cleaning procedures, with a varying percentage of rejected trials. Approximately 75% of the trials were rejected in the multi-level data cleaning, while 53% were rejected in the amplitude-based procedure. However, the analysis findings did not differ between preprocessing pipelines, and the authors conclude that both manual multi-step and automated single-step pipelines were equivalent. Subsequent studies from the Budapest group only use the automated amplitude-based rejection step.

Kujala et al. [60] differed in their approach by applying a manual inspection of independent component analysis (ICA) components to mitigate muscular and other artifacts. Artifact-related components were visually identified and excluded, while a general linear model was used to remove potential electric leakage from the stimulus trigger signal to EEG channels.

In contrast, Levitt and colleagues [66] trained a support vector machine (SVM) classifier for the automated detection of EEG artifacts in human, canine, and rodent subjects. The models showed relatively high accuracy across species in identifying artifacts caused by skeletal and ocular muscles, with an accuracy of 80.57% and an AUC-ROC value of 0.87 for canines specifically.

### Wakefulness EEG in Dogs

Awake dog EEG activity accounts for eight of the 22 examined studies in this revision. One of the studies, Levitt et al., focused on training a SVM model to detect artifacts for preprocessing. Two of the remaining seven studies were exploratory, with Kujala and colleagues [61] showing for the first time the ability to perform non-invasive EEG in eight awake dogs in 2013. They observed changes in the power spectrum over the P3/P4 (parieto-occipital) electrodes during the presentation of a visual stimulus vs rest. Karpinski et. al. [54] recorded pilot data from two companion dogs during rest and after two different commands, also observing qualitative changes in the power spectrum over the F3/F4 (frontal) electrodes.

The remaining five studies [10,17,60,70,99] deployed event-related potentials (ERP) frameworks to understand visual, auditory and language processing. Törnqvist et al. [99] investigated the ERPs of dogs in response to human and dog faces. The study found that ERPs corresponding to early visual processing were detectable at 75–100 ms from stimulus onset, and significant differences for dog and human faces could be identified at around 75 ms at posterior sensors. Another study [60] deployed a similar experimental paradigm, with the addition of emotionally valenced faces and objects, and detected a group-level response sensitive to emotional expressions at 130–170 ms, and the differentiation of faces from objects occurring at 120–130 ms. The authors also trained a support vector machine (SVM) classifier on the EEG data to discriminate between responses to pairs of images. The classification accuracy was highest for humans/dogs vs. scrambled images, with the most informative time interval being between 100–140 ms and 240–280 ms after the presentation of stimuli.

Two studies investigated facets of language processing using ERP paradigms [10, 70]. Magyari et. al [70] explored the group-level ERPs of dogs listening to known, unknown, and nonsense words, finding a significant difference in ERP values between known and nonsense words at 650–800 ms. They also found a positive association between the word usage frequency by the dog’s guardian and the individual dog’s ERP effects, providing evidence that the association may arise from familiarity with the words. The second study to deploy an ERP paradigm [10] investigated the neural processes underlying speech segmentation. The authors examined ERPs from the presentation of artificial words, after participants had been exposed to a continuous speech stream that differed in the distribution of words. In their study, two important factors were considered: transitional probability and word frequency. Transitional probability refers to the likelihood of one sound or word following another in a sequence, reflecting the statistical regularities of the language. Word frequency, on the other hand, represents how often a word occurs in a given language. The results of the study showed that the ERPs exhibited an early effect (220-470 ms) related to transitional probability and a late component (590-790 ms) modulated by both word frequency and transitional probability, indicating the involvement of multiple cognitive processes in speech segmentation.

Finally, Balint et al. [17] investigated the auditory processing of 17 dogs in response to human and dog vocalizations. They found that, similar to humans, dogs exhibited differential ERP responses based on the species of the vocalizer. Specifically, within the time window of 250-650 ms after stimulus onset, ERPs were more positive for human vocalizations compared to dog vocalizations. Furthermore, a later time window of 800-900 ms demonstrated an ERP response that also reflected the species of the vocalizer. These results highlight the existence of species-specific processing of vocalizations in dogs and provide insights into the neural mechanisms underlying their perception of human and non-human vocalizations.

### Sleep EEG in Dogs

EEG recordings of sleeping dogs were examined in 14 of the 22 studies described in this review. In 2014, Kis and colleagues pioneered the use of non-invasive EEG to record brain activity in sleeping dogs, providing a tool to investigate fundamental questions about sleep architecture in those [56]. While most studies relied on human coding and analysis, automated techniques using algorithms to find specific patterns called spindles [43, 45] were developed and refined. Furthermore, machine learning models, including logistic regression (LogReg), gradient boosting trees (GBTs), as well as convolutional neural networks (CNNs), were also deployed and validated to predict sleep stages in dogs [31].

Another study investigated the impact of pre-sleep activity, timing, and location on sleep macro-structure, such as the duration of sleep and the transitions between sleep stages [16]. The authors discovered that the intensity of presleep activity and the location and timing of sleep sessions had interactive effects on sleep macrostructure. Pre-sleep intensive activity and night-time sleeping were associated with more time spent in both non-rapid eye movement (NREM) and rapid eye movement (REM) sleep. Furthermore, they found that dogs sleeping in a location outside their home were less likely to experience REM sleep. A later study by Reicher and colleagues [92] investigated the well-known first-night adaptation effect seen in humans and found that it also manifests in dogs, albeit with marked differences. The first-night adaptation effect refers to the recurring observation that the first recorded sleep session in humans differs from all subsequent recordings, as it is marked by the necessity to adapt to the recording conditions. In dogs, a significant difference was observed between the first and third recordings, with dogs spending more time in sleep and having a shorter latency to drowsiness in session 3 than in session 1. Reicher and colleagues [93] also investigated whether dogs exhibit functional hemispheric asymmetry, a phenomenon in which the right and left hemispheres of an individual displays differential activity during a cognitive process, frequently observed during sleep in aquatic mammals [72]. They found a complex asymmetry contingent on the recording session, sleep cycle, and type of frequency, with some similarities but also many differences between canines and humans.

In addition to exploring fundamental and comparative questions, researchers have also investigated the impact of biological variables such as age, sex, and weight on sleep activity. The impact of age on sleep macrostructure is significant, showing correlations with the power of some frequency bands [91]. Specifically, past 8 months of age, older dogs had higher powers of alpha, beta and gamma frequencies, and lower delta frequencies, compared to younger dogs. Another approach has involved the measurement of spindles, which are phasic bursts of thalamo-cortical activity that appear in the cortex as transient oscillations in the sigma range (typically defined in humans as 9-16 Hz) [21]. In 2017, Iotchev and colleagues [43] developed an algorithm to quantify sleep spindles in dogs and subsequent work has discovered associations between the frequency, density, and amplitude of spindles with the age and sex of canine participants. It is worth highlighting the observation that an increase in age was associated with a decrease in the density and amplitude of slow spindles [44].

Another theme explored by researchers in dog EEG is the relationship between sleep activity and other cognitive processes. For example, Kis and colleagues [55] observed a connection between sleep activity and learning rates, specifically that increased beta and decreased delta activity during REM sleep were related to higher performance on a novel learning task. Which could be related to the processed described on Iotchev’s work, on the potential significance of spindle activity for learning and memory processes [43,45,46]. These studies suggest that learning gain (increases in performance on cognitive tasks between sessions inter-spaced by sleep) is correlated with measures of spindle density. While the authors acknowledged potential confounders with demographic variables, the studies suggest a possible causal role of spindles in the consolidation of memory.

On a different association with sleep quality, Carreiro et al. [19] investigated the relation between sleep activity and owner-rated hyperactivity and found that dogs rated as more hyperactive and impulsive demonstrated less total sleep time, a reduced percentage of REM sleep, and lower spindle density compared to dogs rated as less hyperactive and impulsive. Moreover, owner-rated hyperactivity and impulsivity were associated with increased wakefulness after sleep onset and greater sleep fragmentation.

Finally, two studies explored the relation between sleep activity and features of human-canine interaction - specifically cooperation and attachment. Kiss et al. [57] deployed an experimental paradigm testing the ‘audience effect’ between dogs and their human guardians, which relates to the difference in task performance based on the presence of visual attention. Spectral sleep analysis revealed associations between REM and non-REM power activity and susceptibility to the audience effect. In other words, the willingness of participant dogs to follow task instructions, irrespective of whether their guardian was looking at them, was associated in a trait-like manner with alpha, beta, theta, delta frequency bands power during sleep. Carreiro et al. [18] used an adapted form of the Strange Situation Task (SST) [2]

to index attachment levels of canine subjects and investigated whether derived attachment scores correlated with sleep activity features. They found associations between the level of attachment and the duration of NREM sleep, as the activity in certain frequency bands.

## Discussion

### Review of Findings

### Temporal nature of dog cognitive processes in wakefulness

Non-invasive EEG uses electrodes placed on the scalp that pick up signals that are the end product of the integration of postsynaptic potentials of hundreds of thousands of neurons traversing from the brain across tissue, bone, muscle, skin and hair. This leads to a measure of brain activity with a low spatial resolution but high temporal resolution [22]. Effectively, this means that the inferences from awake canine EEG data are related to fine-grained features of temporal activity. This is what we saw from the five studies that deployed hypothesis-driven ERP analysis frameworks to investigating facial, vocalization and speech comprehension, with the significant time-windows displayed in Table 4. Meaningful inferences can be derived from a wide range of times (between 30-950ms) post onset of a stimulus, as previously highlighted by a study on the potential of machine learning (ML) models such as Support Vector Machines (SVMs) in predicting stimulus categories based on activity in such time-windows [60].

**Table 4:** Overview of selected time-windows that had significant event-related potential activity.

| Time-window (ms) | Description | Studies |
| --- | --- | --- |
| 30-40 | Aggressive dog faces | [60] |
| 75-110 | Difference between faces/objects and scrambled images | [60, 61] |
| 220-470 | Word segmentation (transitional probability) | [10] |
| 250-650 | Species vocalization sensitivity | [17] |
| 360-400 | Difference between dog and human faces | [60] |
| 590-790 | Word frequency | [10] |
| 650-850 | Known versus nonsense words | [70] |
| 800-900 | Valence in vocalizations | [17] |

### Relationship between sleep and physiological traits in dogs

In contrast to wakefulness dog studies, the reviewed sleep studies took a different approach in their analysis framework. While the high temporal resolution of EEG was occasionally leveraged in the spindle studies [44, 45], the main focus was on investigating associations between more general sleep stages and patterns and psychological traits. Some of the significant correlations found are highlighted in Table 5.

**Table 5:** Overview of selected associations between sleep macrostructure and patterns of sleep and physiological and psychological states.

| Sleep feature | Associated physiological or psychology state/trait | Studies |
| --- | --- | --- |
| Duration in NREM and REM | Pre-sleep intensive activity, time and location of sleep, owner-rated hyperactivity | [16, 19] |
| Alpha frequency power | Age, cooperation, attachment | [18, 57] |
| Beta and delta frequency power | Age, learning rate, cooperation | [55, 57] |
| Gamma frequency power | Age | [92] |
| Frequency, density, and amplitude of spindles | Age, sex, learning gain, owner-rated hyperactivity | [43-46] |

### Comparisons Between Dog and Human EEG

Non-invasive EEG with humans has a rich literature, and reviewed studies often used comparative framings to generate hypotheses or provide explanatory models. An overview of some of the overlapping components are provided in Table 6. The N1 component, well-studied in human subjects [11], appears to also be present in canines. Kujala (2013) [61] noted a deflection at 75ms in response to visual stimuli, earlier than typically observed in human studies. This component, observed primarily in posterior channels (P3/P4), appears to differentiate between human and dog faces. Boros (2021) [10] observed a N100 effect at electrode Fz as opposed to electrode Cz for word stimuli. A face-sensitive component, representing a holistic representation of a face, appears at 170ms post stimulus for humans [108]. Kujala (2020) [60] identified emotional expression-dependent effects between 127–170ms from stimulus onset, suggesting face processing in dogs may be connected with the processing of the affective content of the stimulus. The word-familiarity effect, visible in human infants between 200-400ms [75] post word-onset, may also be present in dogs. ERP differences between WORDS and NONSENSE conditions appeared between 650-800ms post word-onset, towards the end of the words [70].

**Table 6:** Overview of selected overlapping EEG components between humans and dogs.

| Component | Human | Canine |
| --- | --- | --- |
| N1 (100ms) | First negative deflection post-stimulus onset | Occurs at 75ms in response to visual stimuli, can differentiate between human and dog faces. |
| Face-sensitive (170ms) | Holistic representation of a face is generated at 170 ms post stimulus onset | Emotional expression-dependent effects at 127–170 ms post stimulus onset, suggesting face processing in dogs may be connected to the processing of the affective content |
| Word-familiarity (200-500ms) | Observable word-familiarity effect in human infants | ERP differences between WORDS and NON-SENSE conditions appeared between 650 and 800 ms following word-onset |
| P300 and LPP (300ms) | Associated with attention and stimulus evaluation | Dogs show a differential ERP response depending on the species of the caller between 250 and 650 ms, suggesting attentional differences to human and dog vocalizations |
| N400 (400ms) | Indicates speech segmentation of candidate words | Significant effect of transitional probability (220-470 ms) and possible higher order recognition (590-790 ms) after word onset |

The P300 is a component of the event-related potential (ERP) that is typically elicited in the process of decision making [97]. It is most commonly evoked during “odd-ball” tasks, where the participant is asked to respond to infrequent or unexpected stimuli amongst a set of standard stimuli. The P300 wave occurs roughly in the range 300-600 milliseconds after the presentation of the stimulus and is understood to be correlated with the degree of attention allocated to a stimulus [87]. Dogs exhibited a P300 response in the study by Balint (2022) [17], with a more positive ERP response to human as compared to dog vocalizations between 250 and 650 ms, which may reflect a difference in the motivational significance and allocated attention to human and dog vocalizations.

Lastly, the N400 component is a well-researched event-related potential (ERP) [62, 63] that is often associated with semantic processing in language comprehension. It’s characterized by a negative peak occurring around 400ms post-stimulus onset. Dogs exhibited a more positive deflection for words with high transitional probability than with low transitional probability between 220-470 ms after word onset, with late ERP effects also observed between 590-790 ms, potentially representing a higher order recognition at the word level.

### Avenues for future development Unexplored questions

While reviewed studies explored the neural correlates of visual and auditory processing, olfactory processing has so far been unexplored. Studies with humans indicate that relevant olfactory processing features can be extracted from EEG data [27, 41]. An understanding of the neural correlates of olfaction would be vital to a greater understanding of a dog’s perception and cognition. Early studies that measured EEG of sedated dogs showed promise in decipherable differences between evoked potentials between stimuli [40] and multiple studies using fMRI have observed meaningful neural correlates from olfactory tasks in dogs [50, 89, 90].

Similarly, studies exploring questions of cognitive control or executive function, defined by Gazzaniga and colleagues as the “set of psychological processes that enable us to use our perceptions, knowledge, and goals to bias the selection of action and thoughts from a multitude of possibilities” [30], have remained relatively underexplored. Cognitive control encompasses a wide range of processes, including working memory, attentional control, cognitive flexibility, and inhibitory control, and some precedent for into these domains is provided by prior canine cognitive research using fMRI [24, 64]. As with olfaction, questions of cognitive control not only inform us about the neural underpinnings of canine cognition but also have significant practical value in the way humans communicate with and train dogs.

Intertwined with these questions on cognitive control are questions on the onset and processing of emotions. Both Kujala et. al. [60] and Balint et. al. [17] investigated the effect of positive and negative stimuli, and found observable differences in the respective evoked potentials. A broader and deeper investigation into the realm of canine emotions would likely elucidate further such evidence, as work with fMRI on emotions such as jealousy shows [23, 53]. Relatedly, it is worth investigating whether fundamental physiological states, such as hunger, stress, and the need of elimination, are also represented by neural correlates that can be be consistently identified by non-invasive EEG. A fundamental constraint on such questions is the poor spatial depth of EEG, as emotional and physiological processing involve structures located deeper in the brain, to which EEG is not the adequate technique to map.

Another underlying challenge with some of these questions is the problem of ascertaining ground truth. This is especially made apparent by the question of identifying emotions and to create ethical experimental conditions where specific emotions can be consistently engendered in a non-human animal. However, there is potential in cross-modality methods being used to triangulate such ground truth, such as the use of machine vision [9] and FACS (Facial Action Coding System) [102] as well as physiological data on respiration and heart-rate from wearable sensors [13, 28].

Finally, the majority of reviewed studies deployed group-level analysis frameworks. Such frameworks have dominated canine cognition research, indeed Arden and colleagues find that from 1911 to 2016, only three studies took an explicit individual-differences approach to exploring canine cognition [6]. Given the large inter-species variation amongst dog breeds, especially in head and brain shape, a greater number of individual-differences analyses is well-warranted.

### Standard Setups for Dog EEG

A challenge downstream of canine variance is the difficulty in standardizing electrode montages. The convention used in human research is the 10-20 system that allows for a consistent placement of electrodes across individuals [49]. Reviewed studies borrowed from the human 10-20 system in the placement of electrodes, although they differed in their labels for similar electrode montages. An important question is if the 10-20 system is capable of transferring over to canines in a useful way given the marked and asymmetrical difference in head shapes between dolichocephalic, mesocephalic, and brachycephalic dogs [96]. For instance, the distance of a 10% posterior increment between a greyhound and a bulldog would likely lead to stark differences in the brain regions that are recorded by the same electrode position [37].

A related issue is the lack of a standard reference electrode. Three different electrode position labels were used by the six different setups, and it is not clear that these labels refer to the same anatomical position. As mentioned prior, the position of the reference electrode has a strong and irrevocable influence on the EEG recording, and it is unlikely that meaningful comparisons can be made across subjects and labs, if the same anatomical reference is not used. Four studies used the ear as a reference position, which runs contrary to the 2020 recommendations for reproducible human EEG research issued by the Organization for Human Brain Mapping (OHBM) [85], which recommended against physically linked earlobe or mastoid electrodes as they are not a neutral reference and can introduce distortions in the data that make modelling intractable. This is likely the case for canine EEG as well, especially given the natural tendency for dogs to move their ears to attune to stimuli, although it is important to note the influence of breed type, as Beagles arguably display less ear movement given their floppy nature. The other four setups either use Cz or Pz as a reference. It is worth noting, as the authors themselves do [17], that the choice of Pz could lead to an attenuated recording from Cz relative to Fz, given the relative distance between the two. Future research could incorporate other reference systems such as bipolar montages, where each channel represents the potential difference between two adjacent electrodes, or the laplacian montage, where the reference is averaged signal of neighboring electrodes [1].

The development of a standard montage and validation of different reference montage systems, centered on the specificity of canine anatomy, and equipped to deal with the large variance amongst canine individuals would be greatly beneficial to future progress in the field.

### A Standard Data Structure for Dog EEG

As the volume and complexity of cognitive neuroscience methods has grown, several challenges emerged in the organization, dissemination, and analysis of data, leading to the creation of new standards and protocols for neuroscience data structure and management. The brain imaging data structure (BIDS), first proposed in 2016 for magnetic resonance imaging [33], is an exemplar of such a standard that embodies the FAIR principles of findability, accessibility, interoperability, and reusability [104]. Recently, a BIDS standard for EEG data - EEG-BIDS - was proposed to address the same concerns [86]. BIDS allows for the friction-less sharing of data within and between laboratories, as well as enabling the automation of analysis scripts, that all serve to address the replicability of findings.

As noted prior, none of the reviewed studies had datasets that were fully accessible to external researchers. Three studies that did attempt to make data open-source lacked the necessary information to replicate analysis, either omitting raw data or lacking vital meta-data. While researchers should be continued to be encouraged to open-source their data, in the vein of suggested guidelines by the Organization for Human Brain Mapping (OHBM), it is also vital that such shared data is interoperable and accessible for researchers across labs and time. Adapting EEG-BIDS for canines would be a crucial step to ensuring replicable and robust canine EEG research, and can readily be done, with the primary challenge being the adoption of a standard anatomical coordination system to serve as the necessary metadata file. As such a data structure would require adoption by researchers in the field and it should be a priority to ensure consensus on its creation and use. Additional canine specific metadata, such as head size measurements, might also be included in such a protocol to allow for flexibility and refinement in analysis. Upon the adoption of canine EEG-BIDS, data can be made accessible on a neuroscience-specific repository such as OpenNeuro, which would allow for large datasets. Furthermore, common analysis pipelines, such as developed by the BCI2000 open source system for humans [94], could be readily adapted to canine data, opening up the world of canine brain-computer interfaces.

### Improving Signal-to-Noise Ratio

The presence of a furry and muscular scalp makes improving the Signal-to-Noise Ratio (SNR) an important challenge to overcome for non-invasive canine EEG experiments. One approach to improving SNR would be using electromyography (EMG), alongside EEG, to quantify the contribution of scalp muscle activity. With enough measurements from individuals and across dog breeds, it could be possible to regress out muscle activity from neural activity, and allow robust recordings from further electrode positions.

Alongside, impedance benchmarks for different breeds, as well as for different electrode types (e.g. wet vs dry), would inform optimal electrode design to increase SNR. An example of such a study was performed by Luca and colleagues for wired vs wireless sub-dermal electrodes with canines [67]. The use of custom canine phantoms to measure impedance for different systems could also be productive, as seen for human EEG studies [35, 58, 81]. It is worth noting that the majority of reviewed studies used wired systems and only one study used a canine-specific cap to hold electrodes in place. The use and development of canine-specific caps coupled with wireless modern systems would greatly reduce the noise from electrode slippage and wire movement [79] and lead to an increase in SNR. Moreover, such systems would allow experiments in freely moving dogs in naturalistic settings.

Along with custom wireless canine systems, the use of active electrodes, over the passive one used by all studies, have the potential to greatly boost SNR. Active electrodes consist of electrodes with a mini-amplifier system loaded onto the electrode itself, boosting signal quality at the source, and thus increasing SNR overall [34]. Another avenue could be the creation of ground-truth standardization tests based on steady-state visual-evoked potentials (SSVEPs) [4, 80]. A SSVEP is the brain’s evoked potential in response to the presentation of periodically flashing visual stimuli, and this has been observed to form a stable EEG wave with a frequency that matches the presented stimulus. Thus, a SSVEP framework can provide a grounding upon which the SNR of a particular system can be measured. Auditory steady-state responses (ASSRs) are similar to SSVEPs but with specific frequencies embedded in audio. ASSR paradigms are potentially easier to do with dogs than SSVEPs as dogs don’t have to be trained to stare at a flashing stimulus. Similar to SSVEPs, ASSRs can provide a metric to ascertain the SNR of a system [77].

Finally, the detection of artifacts is crucial to ensuring high SNR. All awake experimental reviewed studies deployed manual or algorithmic preprocessing pipelines to clean their recorded canine EEG data. However, as shown by Levitt and colleagues [66], machine learning models are capable of eye-blink and muscular artifact identification and removal for non-invasive canine EEG data, although it is important to point out the limited accuracy of 80%. Moreover, the same models used for human EEG data performed well when trained on canine EEG data, suggesting the potential for transferring models with weights from human EEG data to canine EEG data. While the use of ML artifact removal pipelines continues to be debated in the field [47], and further larger studies should be conducted before making any strong conclusions, it seems possible for ML preprocessing pipelines to be a valuable addition to the field, especially as more than half the data from some awake dog studies had to be manually thrown out because of artifacts [70]. As such pipelines could feasibly allow the identification and removal of artifacts without losing out on the data from the entire trial, this could significantly increase the quantity of data available for analysis, boosting SNR, especially as the number and quantity of EEG recordings increases.

### A search for better models for canine EEG

We observed in the reviewed studies a shift in focus from explanatory models to prediction models, a distinction articulated by Yarkoni and Westfall in their 2017 paper [107]. The distinction is raised with the pertinent criticism that psychology often prioritizes explanatory models, which are prone to overfitting and rarely tested for out-of-sample accuracy. They further argue that this emphasis on explanatory models has contributed significantly to the replication crisis in psychology, as these models often fail to generalize to different data sets. In contrast, a predictive approach aims to construct models that can effectively predict out-of-sample data by leveraging established machine learning techniques such as leave one-out cross-validation [12, 107]. This shift toward prediction models offers a promising direction for improving the robustness and generalizability of canine EEG research.

A particular pitfall to be addressed, is the observed limitation in some reviewed studies, where multiple comparisons went uncorrected or under-corrected. As the nature of EEG studies involves many degrees of freedom, including multiple channel locations, time windows, and subjects, spurious significance values are likely. Some studies attempted to adjust for this using methods such as the Bonferroni correction, which involves adjusting the *α* value by the number of hypotheses (m) by the following formulae — actual alpha = desired *α*/m. However, some studies omitted to consider the true range of hypotheses being tested, e.g. not considering time-window choice as a relevant hypothesis. This lack of adequate correction for multiple comparisons, coupled with the relatively small sample size of studies raises serious questions of replicability.

One challenge with the use of predictive models is the need for large amounts of data to train models [26]. One tractable approach could be to combine a predictive model framework with an individual level analysis. That is, large amounts of EEG data can be collected from a few dogs for a specific task, and machine learning models can be trained on data from these individual dogs. Comparing the performance of such models across dogs and tasks, as well as correlation between model performance with other behavioral and cognitive traits should provide insight on questions of canine cognition, whilst providing an alternative model to solve the problem of multiple comparisons.

## Conclusion

The rise of non-invasive canine EEG can be traced to lying at the intersection of three trends — the increasing maturity of cognitive neuroscience, the rejuvenation of canine science, and the increasing sophistication in portable and accessible neuroimaging methods. The latter is primarily due to the increasing interest in brain-computer interfaces (BCIs) [101]. We may be at the cusp of the emerging field of canine brain-computer interfaces, where wearable and non-invasive systems could allow dogs to interact with objects, their environment, and humans through cognitive processes alone. Such systems could be beneficial to researchers in the field of canine science, as it extends the field of possibilities in experimental design, as well as potentially reducing the time needed for operant training. The development of canine BCIs would be benefited by the framings of the field of Animal-Computer Interaction (ACI), which works to design technologies for non-human users using a development model which incorporates iterative prototyping, animal welfare as a central value, and a direct involvement of animal experts at all stages of the development process [100].

It would be vital to remember that neuroscience needs behavior [59], and that further work with canine EEG continues to incorporate behavioral measures alongside neural data. One useful approach could be the lens of embodied and 4E cognition, where cognition is seen to extend beyond the confines of the brain [78,82]. An embodied approach to canine EEG would emphasize embodied data, such as from wearable heart and respiration sensors, as well as acknowledge the influence of the environment, as well as human and con-specifics, on neural patterns.

In conclusion, the utility of non-invasive EEG encompasses the diverse and expansive roles that dogs have come to occupy in our societies, providing a portable, accessible, and ethical method to derive quantitative data on canine cognition. Non-invasive EEG can lead to insights on the shared neurological conditions as well as cognitive processes in humans and canines, and provide a data-driven amplifier to the training and deployment of working dogs.

## Author contributions statement

A.K. and A.Z. conceived the study, A.K. and D.L. conducted the study, A.K, D.L., A.Z. and L.F. analysed the results. All authors wrote the main manuscript text and reviewed the manuscript.

## Notes

### Competing Interest Statement

The authors have declared no competing interest.

